# *Plasmodium falciparum* hydroxymethylbilane synthase does not house any cosynthase activity within the haem biosynthetic pathway

**DOI:** 10.1101/2021.02.09.429727

**Authors:** Alan F. Scott, Evelyne Deery, Andrew D. Lawrence, Martin J. Warren

**Affiliations:** School of Biosciences, University of Kent, Canterbury, Kent, CT2 7NJ, United Kingdom; Current address: School of Biochemistry, University of Bristol, Bristol, BS8 1TD, United Kingdom; Quadram Institute Bioscience, Norwich Research Park, Norwich, NR4 7UQ, United Kingdom

**Keywords:** *Plasmodium falciparum*, haem synthesis, uroporphyrinogen III, hydroxymethylbilane, porphobilinogen deaminase

## Abstract

The production of uroporphyrinogen III, the universal progenitor of macrocyclic, modified tetrapyrroles, is produced from aminolaevulinic acid (ALA) by a conserved pathway involving three enzymes: porphobilinogen synthase (PBGS), hydroxymethylbilane synthase (HmbS) and uroporphyrinogen III synthase (UroS). The gene encoding uroporphyrinogen III synthase has not yet been identified in *Plasmodium falciparum* but it has been suggested that this activity is housed inside a bifunctional hybroxymethylbilane synthase (HmbS). In this present study it is demonstrated that *P. falciparum* HmbS does not have uroporphyrinogen III synthase activity. This was demonstrated by the failure of a codon optimised *P. falciparum hemC* gene, encoding HmbS, to compliment a defined *E. coli hemD^-^* mutant (SASZ31) deficient in uroporphyrinogen III synthase activity. Furthermore, HPLC analysis of the oxidsed reaction product from recombinant, purified HmbS showed that only uroporphyrin I could be detected (corresponding to hydroxymethylbilane production). No uroporphyrin III was detected, thus showing that *P. falciparum* HmbS does not have UroS activity and can only catalyse the formation of hydroxymethylbilane from porphobilinogen.

## Introduction

Haem, as an iron-containing porphyrin, is a modified tetrapyrrole that is derived from the starting material 5-aminolevulinic acid (5-ALA) [1]. The construction of the macrocyclic framework of haem is mediated in just three steps [1]. Firstly, two molecules of 5-ALA are condensed to give a pyrrole, porphobilinogen (PBG), in a reaction catalysed by PBG synthase [2, 3]. The next step involves the polymerisation of four pyrrole units (termed A-D) into a linear bilane called hydroxymethylbilane (HMB) and is mediated by an enzyme called HMB synthase (HmbS) that deaminates and links together four molecules of PBG [4–7]. Finally, the bilane undergoes cyclisation, but only after inversion of the terminal D ring, to give uroporphyrinogen III [6, 7]. These three steps are found in all organisms that make modified tetrapyrroles [1]. These reactions are shown in Figure 1.

**FIGURE 1.**
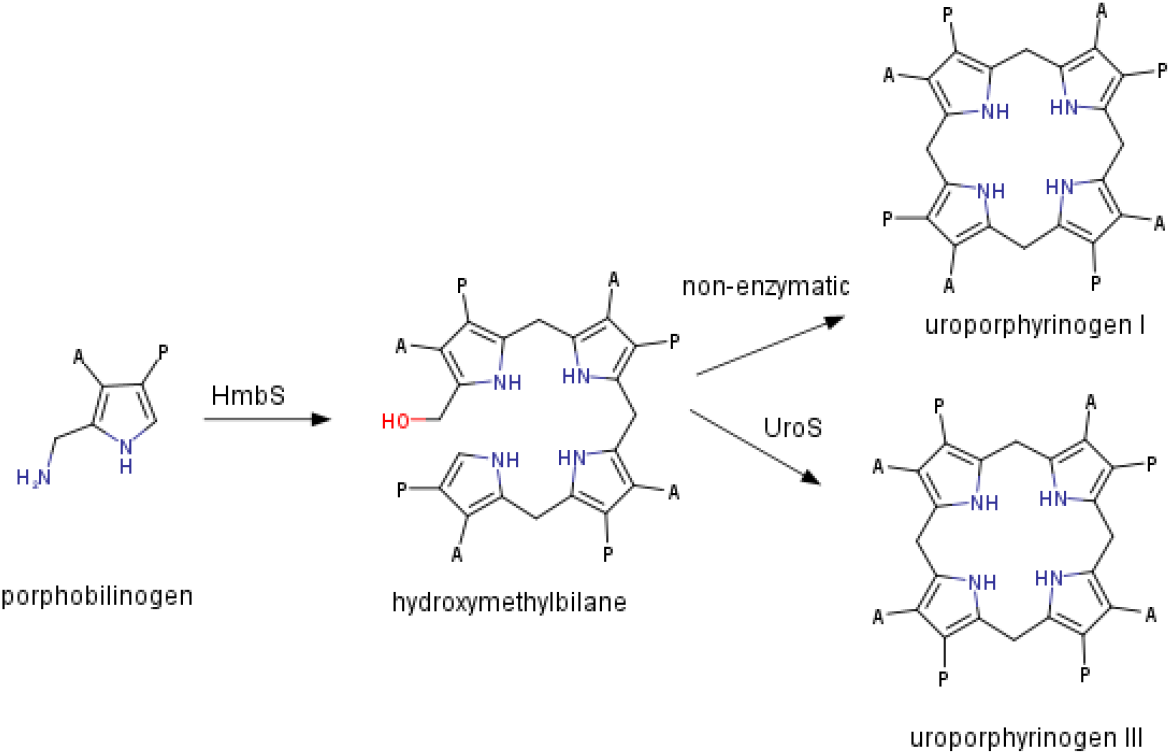
Reactions of HmbS and UroS. HmbS polymerises porphobilinogen into hydroxymethylbilane which auto-cyclises to uroporphyrinogen I. If UroS is present then hydroxymethylbilane is cyclised into uroporphyrinogen III, a reaction that involves the inversion of ringD. A = Acetic Acid, P = propionic acid.

A pathway for haem biosynthesis is found in *Plasmodium falciparum*, the protozoan parasite and causative agent of malaria [8]. As a haematophagous organism (blood-feeding parasite) it exists for part of its life cycle in a haem-rich environment and releases large quantities of haem as an insoluble crystalline material called haemozoin [9]. Nonetheless, for survival outside of the red blood cell it requires a functional haem synthesis pathway that is essential in the liver and parasite growth stages [10, 11]. Recent studies have biochemically characterised the complete set of haem synthesis enzymes from *P. falciparum* with the notable exception of uroporphyrinogen III synthase (UroS, formerly HemD) [12–18]. This enzyme is sometimes called uroporphyrinogen III cosynthase as it often co-purifies with hydroxymethylbilane synthase (HmbS, formerly porphobilinogen deaminase or HemC) and both of these enzymes are required to make uroporphyrinogen III from PBG [5, 19]. HmbS catalyses the synthesis of an unstable linear tetrapyrrole, HMB [4, 6, 7]. This rapidly cyclises into uroporphyrinogen I unless the cosynthase is present to invert the terminal ring and cyclise HMB into uroporphyrinogen III [6, 7, 20]. This is the only isomer that can proceed through the haem synthesis pathway. A candidate gene encoding UroS in *P. falciparum* has been identified by bioinformatics but there have been no biochemical studies to validate the finding [21]. Another report in the literature has suggested that the parasite does not need a separate cosynthase because UroS activity can be found within a bi-functional HmbS that houses both HMB synthase and uroporphyrinogen III cosynthase activities [18]. The evidence presented for this was HPLC identification of the (oxidised) reaction product as uroporphyrin III from both native and recombinant HmbS when incubated with PBG.

Although such dual activity has previously been reported for HmbS from *L. interrogans,* this is a very different protein from *P. falciparum* HmbS, being a fusion of HmbS and UroS enzymes [22]. Conversely, the *P. falciparum* HmbS is clearly not a fusion protein because it has similarity to other HmbS enzymes throughout its entire sequence length (with the exception of the N-terminal apicoplast localisation sequence but dual activity was claimed for a truncated HmbS without this signal sequence) [18]. There are also some short inserts in the *P. falciparum* sequence, but it is unlikely that UroS activity is contained within these inserts because they are not very long – the longest is 31 amino acids. A multiple sequence alignment is shown in Figure S1. It is, therefore, hard to understand how this enzyme could house two very different activities. Consequently, this report investigates more closely the evidence for dual activity using genetic complementation studies and analytical chemistry.

### Experimental Procedures

#### Gene Cloning

A synthetic, codon-adapted *hemC* gene, encoding *P. falciparum* HmbS, was purchased from GeneArt for optimal expression in *Escherishia coli* (Figure S2) and subcloned into a *pET-3a* vector (Novagen) using *NdeI* and *SpeI* restriction sites (the *pET-3a* had been modified to include an *SpeI* site 5’ of the *BamHI* site). Two further constructs were made containing a truncated version of the *hemC* gene to remove a potential signal sequence from the protein product [18]. The truncated gene was obtained by PCR using the following primers:

5’ primer containing *NdeI* site and start codon:
  CAC<u>CATATG</u>GGCATCAAAGATGAAATTATTATCGG
3’ primer containing *SpeI* site and stop codon:
  CTC<u>ACTAGT</u>TATTTATTGTTCAGCAGG

The PCR product was ligated into a *pET-3a* and *pET-14b* (Novagen) use *NdeI* and *SpeI* restriction sites (both vectors had been previously modified to include an *SpeI* site 5’ of the *BamHI* site).

The constructs were sequenced by GATC Biotech to check for the correct insert, reading frame and absence of mutations.

#### Complementation Studies

A defined *hemD^-^* mutant SASZ31(CGSC#: 7153 Coli Genetic Stock Center, Yale University) [23] was transformed with the following plasmids:

|  |  |
| --- | --- |
| pET-3a | Empty vector |
| pET-14b | Empty vector |
| pET-3a *hemC* | *P. falciparum hemC* |
| pET-3a *hemC*<sup>truncated</sup> | *P. falciparum hemC*<sup>truncated</sup> |
| pET-14b *hemD* | *B. megaterium hemD* [24] |
| pET-14b *hemD* | *E. coli hemD* (a kind gift from Prof. Peter Shoolingin-Jordan, Southampton) |

The transformations were plated onto LB-Agar plates with 100 μg/ml ampicillin and 2 % glucose and incubated at 37 °C for 24 hours. They were examined for growth and left for an additional 24 hours after which the colonies were re-streaked onto fresh plates. After incubation at 37 °C for 24 hours the plates were examined for growth.

#### Protein Overproduction and Purification

BL21^STAR^ (DE3) pLysS (Invitrogen) were transformed with the *pET-14b hemC^truncated^* construct and a 1 litre culture of the resulting strain was grown in LB at 37 °C with shaking to an OD_600_ of 0.6. Gene expression was induced for 20 hours at 19 °C by adding 0.4 mM IPTG. Cells were harvested by centrifugation at 4 000 rpm for 15 minutes at 4° C. The pellet was re-suspended in 15 ml re-suspension buffer containing 20mM Tris HCl pH 8.0, 500 mM NaCl, 5 mM imidazole. Cells were lysed on an ice-water slurry by sonication at 60% amplitude for 3 minutes at 30 second intervals. The lysate was spun for 15 minutes at 19 000 rpm and the supernatant loaded onto a Ni^2+^-Sepharose column (GE Healthcare) pre-equilibrated with re-suspension buffer. The column was washed with re-suspension buffer containing 50 mM imidazole and eluted with re-suspension buffer containing 400 mM imidazole. The protein was buffer exchanged with a PD-10 column (GE Healthcare) into 50 mM Tris HCl pH 8.0, 100 mM NaCl.

#### Identification of the Reaction Product

Purified recombinant HmbS was heated to 60 °C for 10 minutes on a heat block prior to the assay to deactivate any contaminating UroS. HmbS (25 μg) was incubated with 200 μM porphobilinogen at 37 °C in 0.1 M Tris HCl pH 8.0. After 1 hour, the reaction was stopped by diluting 10x into 1 M HCl. The reaction product was oxidised by adding 10 μl of a 1 mg/ml benzoquinone in methanol and incubating for 60 minutes. The mixture was run on an HPLC to identify which uroporphyrin isomer was present. Commercial standards of uroporphyrin I and III (Frontier Scientific) were also run to aid identification.

The uroporphyrin I and III isomers were separated on an ACE 5 AQ column, dimensions 250 mm x 4.6 mm, using an Agilent 1100 HPLC system with a flow rate of 1.0 ml/min. The mobile phase was 1 M ammonium acetate pH 5.16 and the organic phase was acetonitrile. A 100 μl sample was injected onto the column (temperature 25 °C) and the porphyrins were detected by their absorbance at 405 nm. A gradient elution was used rising from 13 % to 30 % acetonitrile in 25 minutes and held there for a further 5 minutes. This was adapted from the protocol used by [18].

## RESULTS

### Complementation Studies

To test if *P. falciparum* HmbS harbours UroS activity, complementation studies were performed to see if *P. falciparum hemC* (encoding HmbS) could restore growth to a defined *hemD^-^* mutant (SASZ31) lacking UroS activity [23]. Two *P. falciparum hemC* constructs were used, both of which were codon optimised for expression in *E. coli*. One contained the full-length *hemC* gene in a *pET-3a* vector and the other a truncated *hemC* gene, also in a *pET-3a* vector. The truncation removed a signal sequence known to hinder gene expression in *E. coli* and has been shown not to be essential for activity [18].

The *hemD^-^* mutant SASZ31 was transformed with these constructs and with control plasmids. The controls included an empty *pET-3a* as a negative control and plasmids harbouring known *hemD* genes from *Bacillus megaterium* and *E. coli* as positive controls. As these control genes were in a *pET-14b* plasmid; an empty *pET-14b* was also used as a further control.

The resulting strains were grown on LB-Agar at 37 °C and the size of colonies was noted at 24 and 48 hours. To test for the viability of the colonies after 48 hours they were re-streaked onto a fresh LB-Agar plate and incubated at 37 °C for 24 hours. The plates were examined for colonies.

The control plasmids harbouring known *hemD* genes were able to restore normal growth to the *hemD^-^* mutant. However, the empty vectors and both the *P. falciparum hemC* constructs were unable to restore normal growth. This demonstrates that the *P. falciparum hemC* gene cannot compliment an *E. coli hemD^-^* mutant thus showing that *P. falciparum* HmbS does not have UroS activity. The results are shown in Table 1.

**TABLE 1.**
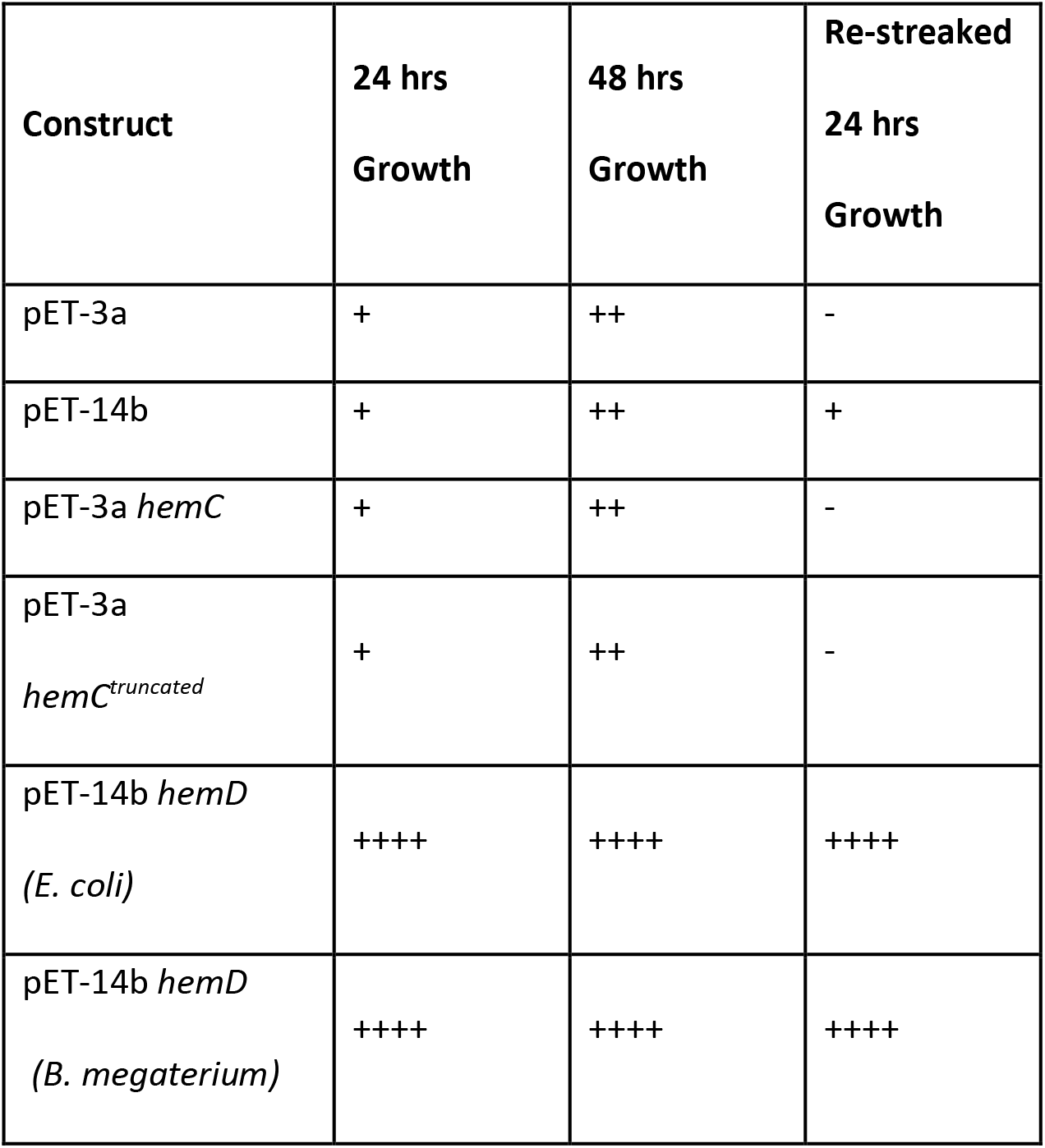
Attempts to compliment a *hemD^-^* mutant with *P. falciparum hemC*. *hemD^-^* mutant SASZ31 was transformed with various constructs and incubated on LB-Agar plates with 0.2 % glucose and appropriate antibiotics at 37 °C for 48 hours. The size of any resultant colonies was recorded after 24 and 48 hours. To test for viability, the colonies were re-streaked onto a fresh plate and grown for a further 24 hours and examined for evidence of growth. The growth is indicated in the table by the number of plus signs from + (poor growth) to ++++ (normal growth). A - indicates that no growth was observed.

| Construct | 24 hrs<br>Growth | 48 hrs<br>Growth | Re-streaked<br>24 hrs<br>Growth |
| --- | --- | --- | --- |
| pET-3a | + | ++ | - |
| pET-14b | + | ++ | + |
| pET-3a *hemC* | + | ++ | - |
| pET-3a<br>*hemC*<sup>truncated</sup> | + | ++ | - |
| pET-14b *hemD*<br>(*E. coli*) | ++++ | ++++ | ++++ |
| pET-14b *hemD*<br>(*B. megaterium*) | ++++ | ++++ | ++++ |

Both the full length and the truncated *hemC* (lacking region encoding signal sequence) *P. falciparum hemC* genes failed to compliment the *E. coli hemD^-^* mutant.

### Protein Overproduction in E. coli and Identification of the Reaction Product

A *pET-14b* construct harbouring the *P. falciparum hemC* gene in frame with an N-terminal hexa-His tag coding sequence was used for protein production in *E. coli*. The *hemC* gene was codon optimised for *E. coli* and lacked the apicoplast localisation sequence. The overproduced protein was mostly insoluble but a small quantity of soluble protein was successfully purified to homogeneity from the cell lysate using Ni^2+^ affinity chromatography. The purity was assessed by SDS-PAGE (Figure S3).

The purified protein was subjected to a 60 °C heat treatment for 10 minutes to deactivate any contaminating UroS. The protein was incubated with substrate for 60 minutes at 37 °C and the resulting product was oxidised with HCl and benzoquinone. This sample was analysed by HPLC to see if the product was uroporphyrin I (corresponding to hydroxymethylbilane) or uroporphyrin III (corresponding to uroporphyrinogen III). Identification was by comparison with commercial standards of uroporphyrin I and III. The product of the reaction product matched the retention time of uroporphyrin I. The results are shown in Figure 2 and show that *P. falciparum* HmbS does not have UroS activity but can only make hydroxymethylbilane.

**FIGURE 2.**
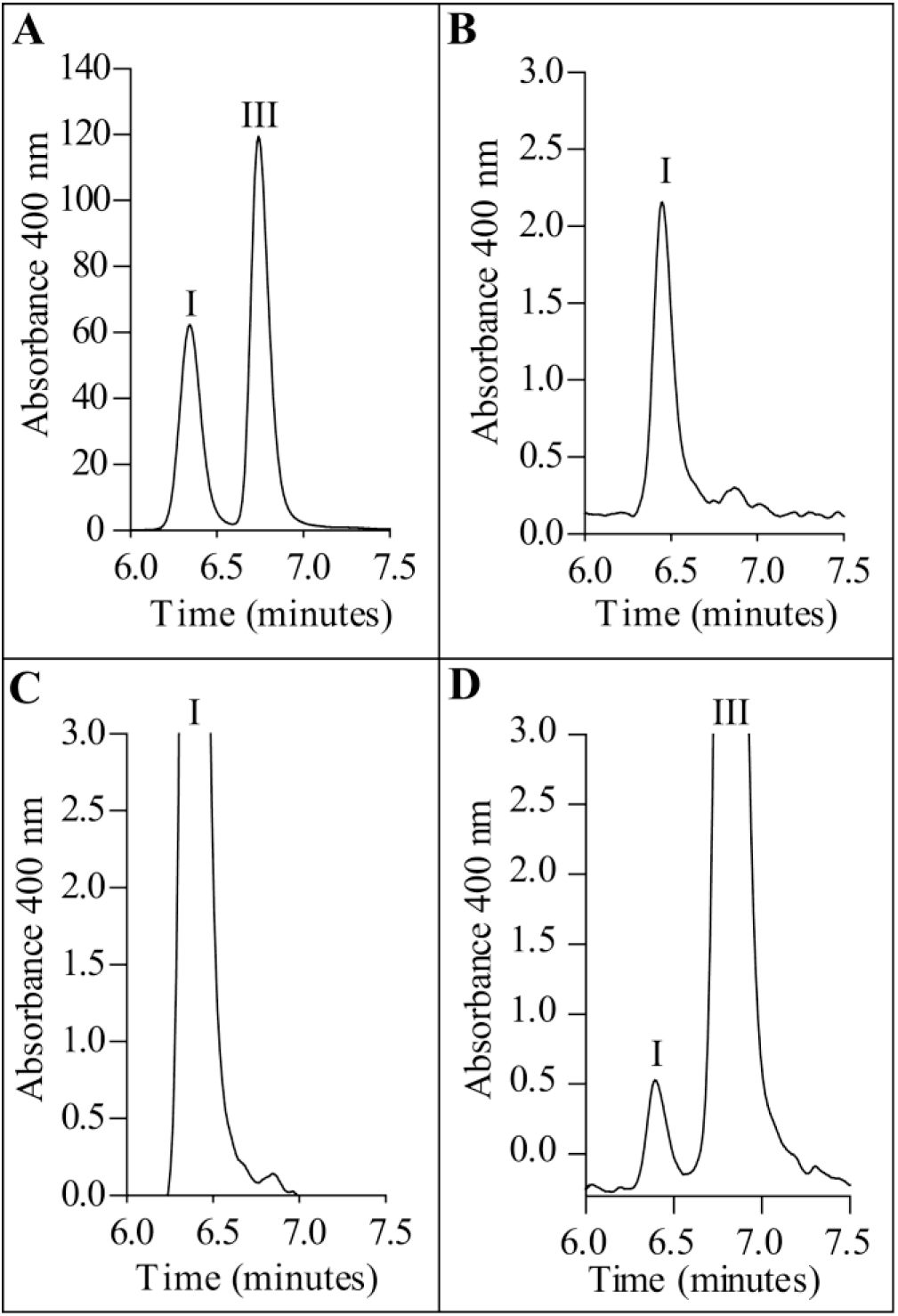
HPLC Analysis of HmbS Reaction Product. HPLC traces showing (A) commercial standards of uroporphyrin I (left) and III (right), (B) the oxidised reaction product of HmbS alone and (C) spiked with uroporphyrin I and (D) uroporphyrin III

## DISCUSSION

The claim that *P. falciparum* HmbS has UroS activity [18] has been challenged through complementation studies with a *hemD^-^* mutant and HPLC analysis of the reaction product from recombinant enzyme. SASZ31 is a defined *hemD^-^* mutant that grows very poorly [23]. Complementation with control *hemD* genes from *Bacillus megaterium* and *E. coli* were able to restore normal growth to the mutant but *P. falciparum hemC* could not. Because HmbS has an apicoplast localisation sequence that hinders expression but is not required for alleged dual activity [18], a truncated gene lacking this sequence was also made. This too failed to complement the mutant.

Furthermore, the truncated HmbS was overproduced in *E. coli* with an N-terminal hexa-His tag and purified. After incubation with substrate for an hour at 37 °C, the sample was oxidised and run on HPLC along with commercial standards of uroporphyrin I and uroporphyrin III. The HPLC result clearly identified the enzyme’s oxidised product as uroporphyrin I. No uroporphyrin III could be detected. These results contradict those previously published [18] where HPLC analysis of the reaction product from native and recombinant HmbS identified the (oxidised) reaction product as uroporphyrin III. This conflict could be explained by the presence of a contaminating UroS in the earlier study. Although the researchers used heat treatment to denature any UroS (HmbS is heat stable but UroS is not), it is possible that any UroS could have refolded and re-activated itself during the 12 hour incubation of heat-treated HmbS with substrate [25–27]. Also, the assay buffer contained additives known to increase the stability of UroS [27]. Our results clearly demonstrate the previous claim, that *P. falciparum* HmbS contains uroporphyrinogen III synthase (UroS) activity, is mistaken [18]. It should now be a matter of importance to find the gene which encodes for the real uroporphyrinogen III synthase.

## Acknowledgements

This work was funded by Pfizer.

## REFERENCES

1. Heinemann IU, Jahn M, Jahn D. The biochemistry of heme biosynthesis. Arch Biochem Biophys 2008. 474:238–251. DOI: 10.1016/J.ABB.2008.02.015.

2. Shemin D, Russell CS. δ-aminolevulinic acid, its role in the biosynthesis of porphyrins and purines. J Am Chem Soc 1953. 75:4873–4874. DOI: 10.1021/ja01115a546.

3. Jaffe EK. The Remarkable Character of Porphobilinogen Synthase.. Acc Chem Res 2016. 49:2509–2517. DOI: 10.1021/acs.accounts.6b00414.

4. Battersby AR, Fookes CJR, Gustafson-Potter KE, Matcham GWJ, McDonald E. Proof by synthesis that unrearranged hydroxymethylbilane is the product from deaminase and the substrate for cosynthetase in the biosynthesis of uro’gen-III. J Chem Soc Chem Commun 1979. 0:1155. DOI: 10.1039/c39790001155.

5. Bogorad L. The enzymatic synthesis of porphyrins from porphobilinogen. I. Uroporphyrin I.. J Biol Chem 1958. 233:501–9.

6. Battersby AR, Fookes CJR, Gustafson-Potter KE, McDonald E, Matcham GWJ. Biosynthesis of porphyries and related macrocycles. Part 18. Proof by spectroscopy and synthesis that unrearranged hydroxymethylbilane is the product from deaminase and the substrate for cosynthetase in the biosynthesis of uroporphyrinogen-III. J Chem Soc Perkin Trans 1 1982. 0:2427. DOI: 10.1039/p19820002427.

7. Battersby AR, Fookes CJR, Gustafson-Potter KE, McDonald E, Matcham GWJ. Biosynthesis of porphyries and related macrocycles. Part 17. Chemical and enzymic transformation of isomeric aminomethylbilanes into uroporphyrinogens: proof that unrearranged bilane is the preferred enzymic substrate and detection of a transient intermediate. J Chem Soc Perkin Trans 1 1982. 0:2413. DOI: 10.1039/p19820002413.

8. Surolia N, Padmanaban G. De novo biosynthesis of heme offers a new chemotherapeutic target in the human malarial parasite. Biochem Biophys Res Commun 1992. 187:744–750. DOI: 10.1016/0006-291X(92)91258-R.

9. Egan TJ. Haemozoin formation. Mol Biochem Parasitol 2008. 157:127–136. DOI: 10.1016/J.MOLBIOPARA.2007.11.005.

10. Ke H, Sigala PA, Miura K, Morrisey JM, Mather MW, Crowley JR, Henderson JP, Goldberg DE, Long CA, Vaidya AB. The heme biosynthesis pathway is essential for Plasmodium falciparum development in mosquito stage but not in blood stages.. J Biol Chem 2014. 289:34827–37. DOI: 10.1074/jbc.M114.615831.

11. Goldberg DE, Sigala PA. Plasmodium heme biosynthesis: To be or not to be essential?. PLOS Pathog 2017. 13:e1006511. DOI: 10.1371/journal.ppat.1006511.

12. Varadharajan S, Dhanasekaran S, Bonday ZQ, Rangarajan PN, Padmanaban G. Involvement of delta-aminolaevulinate synthase encoded by the parasite gene in de novo haem synthesis by Plasmodium falciparum.. Biochem J 2002. 367:321–7. DOI: 10.1042/BJ20020834.

13. Dhanasekaran S, Chandra NR, Chandrasekhar Sagar BK, Rangarajan PN, Padmanaban G. Delta-aminolevulinic acid dehydratase from Plasmodium falciparum: indigenous versus imported.. J Biol Chem 2004. 279:6934–42. DOI: 10.1074/jbc.M311409200.

14. Nagaraj VA, Prasad D, Rangarajan PN, Padmanaban G. Mitochondrial localization of functional ferrochelatase from Plasmodium falciparum. Mol Biochem Parasitol 2009. 168:109–112. DOI: 10.1016/J.MOLBIOPARA.2009.05.008.

15. Nagaraj VA, Arumugam R, Prasad D, Rangarajan PN, Padmanaban G. Protoporphyrinogen IX oxidase from Plasmodium falciparum is anaerobic and is localized to the mitochondrion. Mol Biochem Parasitol 2010. 174:44–52. DOI: 10.1016/J.MOLBIOPARA.2010.06.012.

16. Nagaraj VA, Prasad D, Arumugam R, Rangarajan PN, Padmanaban G. Characterization of coproporphyrinogen III oxidase in Plasmodium falciparum cytosol. Parasitol Int 2010. 59:121–127. DOI: 10.1016/J.PARINT.2009.12.001.

17. Nagaraj VA, Arumugam R, Chandra NR, Prasad D, Rangarajan PN, Padmanaban G. Localisation of Plasmodium falciparum uroporphyrinogen III decarboxylase of the heme-biosynthetic pathway in the apicoplast and characterisation of its catalytic properties. Int J Parasitol 2009. 39:559–568. DOI: 10.1016/J.IJPARA.2008.10.011.

18. Nagaraj VA, Arumugam R, Gopalakrishnan B, Jyothsna YS, Rangarajan PN, Padmanaban G. Unique properties of Plasmodium falciparum porphobilinogen deaminase.. J Biol Chem 2008. 283:437–44. DOI: 10.1074/jbc.M706861200.

19. Shoolingin-Jordan PM. Porphobilinogen deaminase and uroporphyrinogen III synthase: structure, molecular biology, and mechanism.. J Bioenerg Biomembr 1995. 27:181–95.

20. Battersby AR, Fookes CJR, McDonald E, Meegan MJ. Biosynthesis of type-III porphyrins: proof of intact ezymic conversion of the head-to-tail bilane into uro’gen-III by intramolecular rearrangement. J Chem Soc Chem Commun 1978. 0:185. DOI: 10.1039/c39780000185.

21. Mohanty S, Srinivasan N. Identification of “missing” metabolic proteins of Plasmodium falciparum: a bioinformatics approach.. Protein Pept Lett 2009. 16:961–8.

22. Guégan R, Camadro J-M, Saint Girons I, Picardeau M. Leptospira spp. possess a complete haem biosynthetic pathway and are able to use exogenous haem sources. Mol Microbiol 2004. 49:745–754. DOI: 10.1046/j.1365-2958.2003.03589.x.

23. Chartrand P, Tardif D, Saaasarman A. Uroporphyrin- and Coproporphyrin I-accumulating Mutant of Escherichia coli K12. J Gen Microbiol 1979. 110:61–66. DOI: 10.1099/00221287-110-1-61.

24. Raux E, Leech HK, Beck R, Schubert HL, Santander PJ, Roessner CA, Scott AI, Martens JH, Jahn D, Thermes C, Rambach A, Warren MJ. Identification and functional analysis of enzymes required for precorrin-2 dehydrogenation and metal ion insertion in the biosynthesis of sirohaem and cobalamin in Bacillus megaterium.. Biochem J 2003. 370:505–16. DOI: 10.1042/BJ20021443.

25. Jordan PM, Thomas SD, Warren MJ. Purification, crystallization and properties of porphobilinogen deaminase from a recombinant strain of Escherichia coli K12.. Biochem J 1988. 254:427–35.

26. Alwan AF, Mgbeje BI, Jordan PM. Purification and properties of uroporphyrinogen III synthase (co-synthase) from an overproducing recombinant strain of Escherichia coli K-12.. Biochem J 1989. 264:397–402.

27. Omata Y, Sakamoto H, Higashimoto Y, Hayashi S, Noguchi M. Purification and Characterization of Human Uroporphyrinogen III Synthase Expressed in Escherichia coli. J Biochem 2004. 136:211–220. DOI: 10.1093/jb/mvh111.

