## Supporting Information for "*Plasmodium falciparum* hydroxymethylbilane synthase does not house any cosynthase activity within the haem biosynthetic pathway"

**Contents:**

S1. Multiple sequence alignment for *P. falciparum* HmbS

S2. Sequence of synthetic *hemC* gene encoding *P. falciparum* HmbS

S3. SDS-PAGE showing purification of recombinant *P. falciparum* HmbS

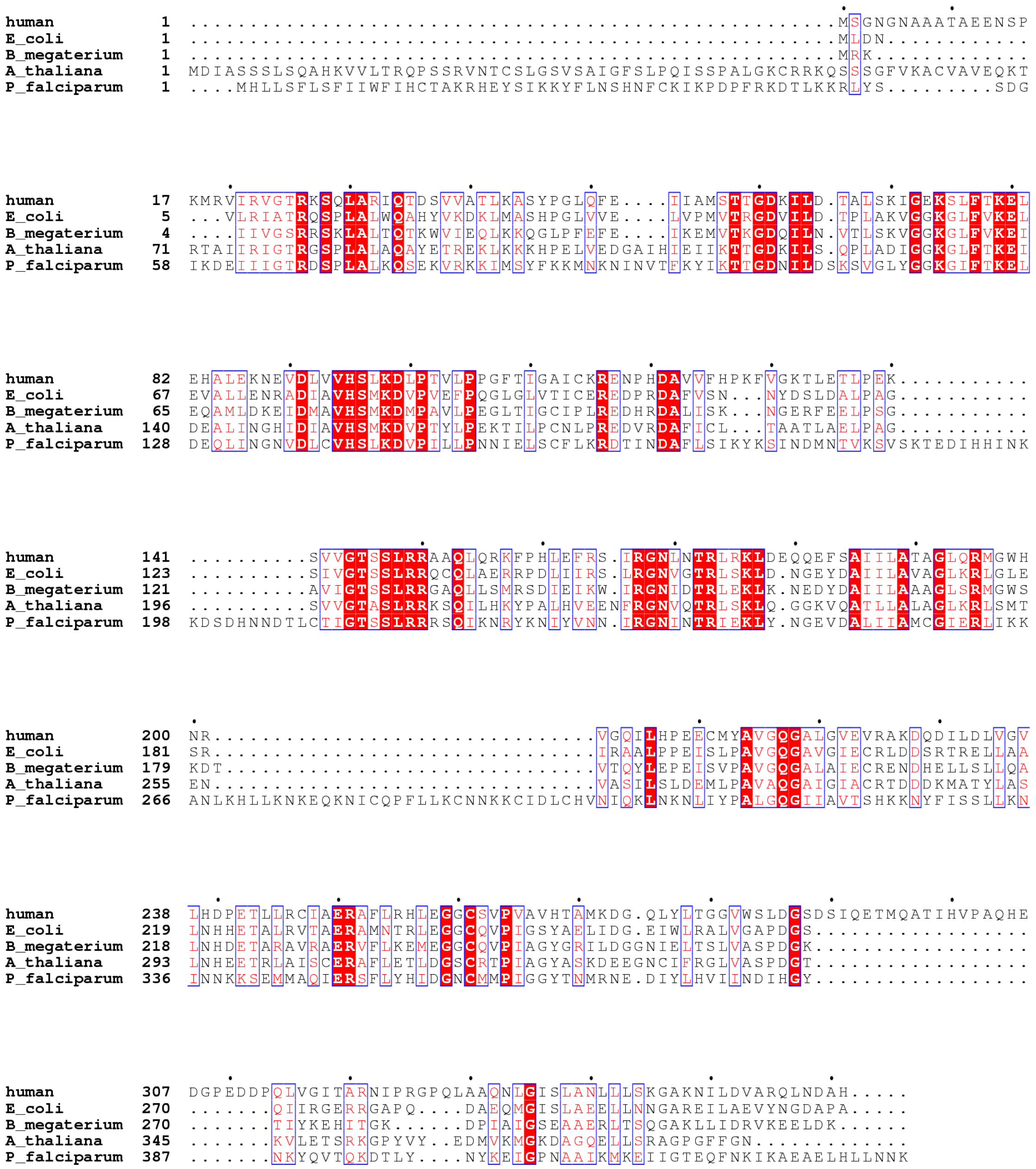


**FIGURE S1 – Multiple sequence alignment for HmbS**

This alignment was produced using ClustalW (http://www.ebi.ac.uk/Tools/msa/clustalw2/) and formatted with ESPript (http://espript.ibcp.fr/ESPript/ESPript/). The HmbS sequences come from Human, *Bacillus megaterium*, *Escherichia coli*, *Arabidopsis thalinana* and *Plasmodium falciparum.*

**ATG**GGCATCAAAGATGAAATTATTATCGGCACCCGTGATAGCCCGCTGGCCCTGAAACAGAGCGAA
AAAGTGCGCAAAAAAATCATGAGCTACTTCAAAAAAATGAACAAAAACATCAACGTGACCTTCAAAT
ACATTAAAACCACCGGCGATAACATTCTGGATAGCAAAAGCGTGGGCCTGTATGGCGGCAAAGGCA
TTTTTACCAAAGAACTGGATGAACAGCTGATTAACGGCAACGTGGATCTGTGCGTGCATAGCCTGAA
AGATGTGCCGATTCTGCTGCCGAACAACATTGAACTGAGCTGCTTTCTGAAACGTGATACCATCAAC
GATGCGTTTCTGAGCATCAAATATAAAAGCATCAACGATATGAACACCGTGAAAAGCGTGAGCAAA
ACCGAAGATATCCACCACATCAATAAAAAAGATAGCGATCACAACAACGATACCCTGTGCACCATTG
GCACCAGCAGCCTGCGTCGTCGTAGCCAGATTAAAAACCGCTATAAAAACATCTATGTGAACAACAT
TCGCGGCAACATTAACACCCGTATCGAAAAACTGTATAACGGCGAAGTGGATGCGCTGATTATTGCC
ATGTGCGGCATTGAACGCCTGATTAAAAAAGCGAACCTGAAACACCTGCTGAAAAACAAAGAACAG
AAAAACATCTGCCAGCCGTTTCTGCTGAAATGCAACAACAAAAAATGCATCGATCTGTGCCATGTGA
ACATTCAGAAACTGAACAAAAACCTGATTTATCCGGCGCTGGGCCAGGGCATTATTGCGGTGACCAG
CCACAAAAAAAACTACTTCATCAGCAGCCTGCTGAAAAATATTAACAACAAAAAAAGCGAAATGATG
GCGCAGATTGAACGTAGCTTTCTGTATCACATTGATGGCAACTGCATGATGCCGATTGGCGGCTATA
CCAACATGCGTAACGAAGATATCTATCTGCACGTGATTATCAACGATATCCACGGCTACAATAAATAT
CAGGTGACCCAGAAAGATACCCTGTATAACTATAAAGAAATTGGCCCGAACGCGGCGATTAAAATG
AAAGAAATCATCGGCACCGAACAGTTCAACAAAATTAAAGCGGAAGCGGAACTGCACCTGCTGAAC
AATAAA**TAA**

**FIGURE S2 – Sequence of synthetic *hemC* gene encoding *P. falciparum* HmbS**

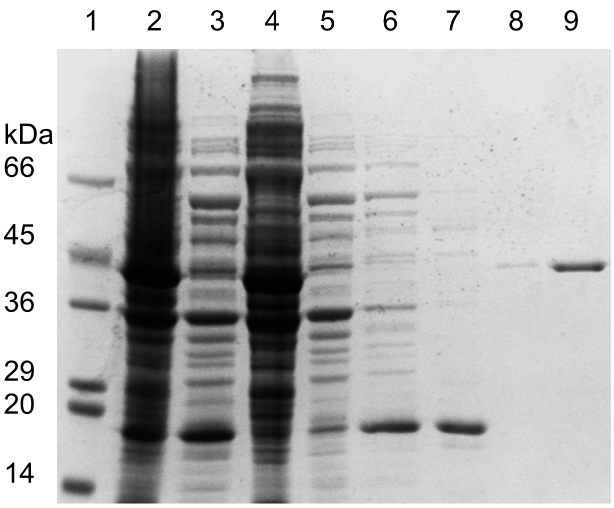


**FIGURE S3 – SDS-PAGE showing purification of recombinant *P. falciparum* HmbS**

(1) Markers SDS7 (Sigma); (2) Total protein from lysate; (3) Total soluble protein from lysate; (4) Total insoluble protein from lysate; (5) Flow through from Ni-sepharose column; (6) Flow through from 5 mM imidazole wash; (7) Flow through from 50 mM imidazole wash; (8-9) Elution fractions from 400 mM imidazole wash

Expected mass of HmbS: 43.6 kDa
